# Dogcatcher2: Improved statistical detection of transcriptional readthrough and repetitive element analysis across sequencing platforms

**DOI:** 10.64898/2026.05.07.723642

**Authors:** Marko Melnick, Christopher D. Link

**Affiliations:** Integrative Physiology, University of Colorado, Boulder, CO; Institute for Behavioral Genetics, University of Colorado, Boulder, CO

## Abstract

Downstream of Gene (DoG) transcription occurs when RNA polymerase II fails to terminate normally at the transcription end site, resulting in extended transcription downstream of the gene. This is a widespread phenomenon linked to cellular stress, cancer and neurodegeneration. Existing tools for DoG detection from short-read RNA-seq rely on absolute coverage thresholds and sliding window approaches that are sensitive to sequencing depth and expression level. Here we present Dogcatcher2, which applies improved statistical detection methods to gene body-normalized coverage profiles. Using long-read ground truth across multiple datasets, we show that Dogcatcher2 outperforms existing methods in both detection sensitivity and boundary accuracy while maintaining high precision even at low sequencing depths. Dogcatcher2 further improves detection on pseudobulk scRNA-seq and snRNA-seq data. Analysis of DoG regions in human reveals specific enrichment for Alu elements including inverted Alu pairs capable of forming double-stranded RNA, with transposable elements within DoG regions showing elevated expression, connecting readthrough transcription to dsRNA generation and innate immune signaling.

## Introduction

Downstream-of-gene transcripts (DoGs) are RNA species generated when RNA polymerase II fails to terminate normally at the transcription end site (TES), resulting in extended transcription of the region downstream of the gene [1,2]. DoGs are induced by diverse cellular stresses including heat shock, osmotic stress, hypoxia, viral infection, and oxidative damage, and have been observed across species from *C. elegans* to human [1,3–6]. DoG expression is also dysregulated in cancer, with differential expression between tumors and matched normal tissues that correlates with patient survival [7]. DoGs can be polyadenylated or non-polyadenylated, are predominantly chromatin-associated, and can extend tens to hundreds of kilobases downstream of the parent gene [1]. Functionally, DoGs that extend into neighboring genes on the opposite strand can generate antisense transcripts with regulatory potential [6].

Several computational tools have been developed for DoG detection from RNA-seq data. DoGFinder was the first dedicated tool, using a binary coverage threshold (requiring 60% of bases covered in each window) with a minimum DoG length of 4 kb [8]. While effective for initial surveys, DoGFinder’s binary coverage criterion is sensitive to sequencing depth, requiring library normalization by downsampling before cross-sample comparison [8]. ARTDeco extended DoG analysis by adding read-in gene detection and readthrough level quantification, using an absolute FPKM threshold (0.15) for DoG boundary calling [9]. Although ARTDeco computes gene body signal ratios, these are calculated as summary statistics after detection rather than being used for boundary calling itself. ARTDeco also reported approximately 5-fold faster runtime than the original Dogcatcher [9]. The original Dogcatcher used a sliding window approach with binary coverage to detect DoGs, PoGs (upstream-of-gene transcripts), and antisense variants (ADoGs, APoGs) in *C. elegans* heat shock data [6].

A critical limitation shared by all existing tools is the use of absolute signal thresholds for DoG detection. Whether using binary coverage (DoGFinder), absolute FPKM (ARTDeco), or coverage fraction (Dogcatcher), these approaches are inherently depth-dependent: the same gene may be called as DoG-positive or DoG-negative depending on sequencing depth alone. None of these tools normalize downstream signal to the parent gene’s expression level during detection. Additionally, no existing tool has been validated against long-read ground truth for DoG boundary accuracy, none have been applied to single-cell or single-nucleus RNA-seq, and no boundary accuracy metrics beyond binary detection have been reported.

Here we present Dogcatcher2, a comprehensive update that addresses these limitations. The key innovation is a suite of five independent statistical detectors (sliding window, PELT changepoint, Bayesian changepoint, segmented regression, and hidden Markov model) from which the user selects one; segmented regression performs best in our benchmarks (see Results) and is the default. These replace the simple sliding window of previous tools. The detector framework is modular, making it straightforward to add new detection algorithms without modifying the core pipeline. These detectors operate on gene body exon-normalized coverage, where downstream signal is expressed as a fraction of the parent gene’s expression before detection occurs, providing expression-level-independent detection. A new long-read mode generates standardized ground truth from PacBio and ONT data with 5’ origin filtering and polyA site filtering. We validate Dogcatcher2 across two multi-platform datasets (SG-NEx and LongBench), demonstrate application to bulk, scRNA, and snRNA data, and provide read-in gene detection with expression correction for differential expression analysis. Dogcatcher2 is implemented in R with minimal dependencies, designed for easy installation and use. The modular architecture including pluggable detectors, configurable filters such as polyA site filtering, and a standardized long-read ground truth pipeline is intended to support ongoing development as the field evolves. We further demonstrate that DoG regions in human are enriched for Alu elements and inverted Alu pairs capable of forming double-stranded RNA, with transposable elements within DoGs showing elevated expression, connecting readthrough transcription to dsRNA generation and innate immune signaling.

## Methods

### Dogcatcher algorithm

The original Dogcatcher was developed for detection of downstream-of-gene transcription (DoGs) in *C. elegans* using a sliding window approach (Melnick et al., 2019). Gene annotation processing, including GTF flattening, nested gene removal, and DoG/PoG/ADoG/APoG classification follows the original implementation. Here we describe significant improvements to detection, normalization, and validation.

#### Gene body exon normalization

A central improvement in Dogcatcher2 is gene body exon normalization. Downstream coverage in each window is divided by a reference signal derived from the parent gene’s own exons before any detection occurs. For DoG detection (downstream), the reference exon set is built by accumulating exons from the 3’ end of the gene until a cumulative length of at least 500 bp is reached. For PoG detection (upstream), exons are accumulated symmetrically from the 5’ end. For genes lacking exon annotations, the last (or first) window-sized region of the gene body is used as a fallback reference. The mean coverage across the reference exon set serves as the denominator for normalization, and the normalized signal for each downstream window is:

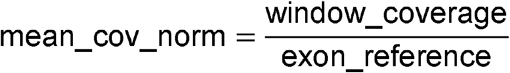

This normalization renders detection independent of absolute expression level. A gene expressed at 10x and a gene expressed at 10,000x will produce the same normalized profile if their readthrough is proportionally equivalent. This is fundamentally different from existing approaches: DoGFinder uses binary base coverage (≥60% of bases covered) which is depth-sensitive [8], and ARTDeco uses an absolute 0.15 FPKM threshold which fails for lowly-expressed genes with genuine readthrough and falsely detects highly-expressed genes with minimal readthrough [9].

Genes must pass a minimum exon signal threshold before detection proceeds. The `min_exon_cov` parameter (default 3x) requires the last exon(s) of the gene (cumulative ≥500 bp) to have at least 3-fold mean coverage; genes below this threshold are skipped entirely. This filter ensures that normalization denominators are meaningful and prevents noise amplification in lowly-expressed genes where stochastic coverage fluctuations would produce unreliable normalized profiles. It replaces the whole-gene-body coverage filter used in the original Dogcatcher, focusing the signal check on the specific exons used for normalization rather than averaging across the entire gene body. In long-read mode, an analogous `min_reads` threshold (default 2) requires a minimum number of readthrough reads per gene before a DoG is reported.

#### Statistical detectors

Dogcatcher2 provides five statistical detectors that all operate on the same exon-normalized coverage profile, allowing users to select the method best suited to their data characteristics and biological question. All detectors scan downstream windows and return a DoG endpoint (or no call if no readthrough is detected).

**Sliding window** extends the approach from the original Dogcatcher [6], walking downstream from the transcription end site (TES) and terminating the DoG when the normalized coverage drops below a threshold (default 5% of gene body signal). This is the simplest detector and provides backwards compatibility.

**PELT changepoint** uses the Pruned Exact Linear Time algorithm [10] to identify the optimal changepoint in the normalized coverage series. This is the point where mean signal shifts from readthrough-level to background. PELT is statistically principled and requires no user-specified threshold; it uses a penalized likelihood criterion to detect the single most significant mean shift.

**Bayesian changepoint** computes the posterior probability of a changepoint at each window position using conjugate normal-inverse-gamma priors. This detector is implemented in pure base R with no package dependencies. The DoG endpoint is placed at the window with highest posterior probability, provided the signal drop exceeds a minimum effect size.

**Segmented regression** fits a piecewise linear model to the normalized coverage profile, identifying the breakpoint where the signal transitions from a decay slope to a flat background. This detector explicitly models the biology of readthrough transcription: coverage decays gradually downstream of the TES rather than maintaining a constant level before dropping abruptly. Our benchmarks show segmented regression achieves the best boundary accuracy among all detectors, with called boundaries closest to the 75th percentile of long-read endpoints.

**Hidden Markov Model (HMM)** fits a two-state model (readthrough vs. background) to the normalized coverage using the Viterbi algorithm. The HMM is best suited for noisy or sparse data where individual window values fluctuate substantially, as it uses transition probabilities to smooth state assignments across adjacent windows.

The detector framework is modular: new detection algorithms can be added without modifying the core pipeline.

#### Read-in gene detection and expression correction

When a DoG extends far enough to overlap a downstream gene, transcription from the upstream gene contributes reads that are counted as expression of the downstream “read-in” gene. This read-in signal does not reflect transcriptional activation of the downstream gene and can inflate expression estimates, generate false differential expression calls, and dilute gene ontology enrichment analyses [9].

Dogcatcher2 identifies read-in genes during DoG detection. For each global DoG (one that reaches 100 kb, the maximum scan distance), the pipeline checks whether the DoG region overlaps a downstream gene on the same strand and reports the downstream gene’s identifier, name, and a log2 read-in ratio calculated as:

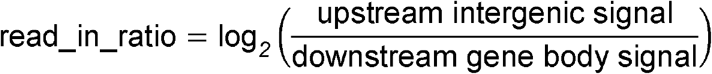

This ratio quantifies the severity of read-in contamination. A high ratio indicates that the upstream DoG signal is substantial relative to the downstream gene’s own expression, suggesting that a large fraction of observed expression may originate from readthrough rather than independent transcription.

For differential expression analysis, Dogcatcher2 provides an expression correction step integrated into the DESeq2 pipeline [11]. After featureCounts quantification and before DESeq2 normalization, the pipeline loads the consensus DoG results, identifies read-in genes, and subtracts the estimated upstream DoG signal from their raw counts:

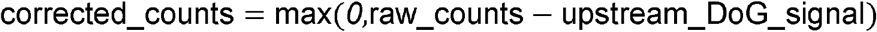

Here, upstream_DoG_signal is the featureCounts read count over the upstream gene’s full DoG interval (TES to detected DoG end). The full DoG count is subtracted from the read-in gene’s gene-body count without modeling how far the readthrough extends into the gene body, which is conservative: if the read-in only contaminates the proximal portion of the downstream gene, this approach over-corrects. We made this design choice because the alternative (scaling the subtraction by an estimated decay profile) would require per-gene long-read data that is not available in most short-read experiments. This correction prevents inflated expression estimates and false DE calls for read-in genes. ARTDeco introduced a similar concept as “gene expression deconvolution” and demonstrated that uncorrected read-in genes produce spurious GO enrichments and eQTLs that actually map to upstream DoG-producing genes [9]. Our implementation integrates this correction directly into the standard featureCounts-to-DESeq2 pipeline.

#### Long-read mode

Dogcatcher2 includes a long-read mode (`--mode longread`) that generates standardized readthrough ground truth from PacBio IsoSeq, PacBio Kinnex, or Oxford Nanopore Technologies (ONT) direct RNA data. Unlike short-read coverage-based detection, each long read represents a single transcript molecule, and the 3’ endpoint of each read directly reports where that transcript terminated. This provides per-read resolution of DoG boundaries rather than aggregate coverage profiles.

For each gene, the long-read mode identifies reads whose 3’ endpoints extend beyond the TES into the downstream intergenic region. A critical 5’ origin filter requires each read’s 5’ end to fall within the annotated gene body, ensuring the read originated from that gene rather than from an overlapping downstream gene that happens to extend upstream. Without this filter, 22-35% of apparent readthrough reads were misattributed from neighboring genes in our datasets. A similar 5’ assignment strategy was used by the LRGASP consortium for read-to-gene attribution [12].

Per-gene output includes: the number of readthrough reads, median and maximum extension past TES, all individual read endpoints, the readthrough fraction (readthrough reads / total gene body reads), and the platform (auto-detected from read length distribution: PacBio uses MAPQ ≥ 20, ONT uses MAPQ ≥ 5). The same flattened GTF used for short-read detection is used for long-read mode, ensuring consistent gene models for fair comparison between detection and ground truth.

### Datasets

#### SG-NEx

The Singapore Nanopore Expression project (Chen et al., 2025) provides matched multi-platform RNA-seq from seven human cell lines. We used the HepG2 hepatocellular carcinoma line: 3 Illumina PE150 replicates, 1 PacBio IsoSeq dataset, and 1 ONT direct RNA dataset, all from the same RNA extraction.

#### LongBench

The LongBench dataset (Ritchie et al., 2025) provides matched bulk and single-cell transcriptomics across eight human lung cancer cell lines. We used three cell lines (H146 SCLC, H1975 NSCLC, H526 SCLC) with the following data per cell line: bulk Illumina PE RNA-seq, PacBio Kinnex long-read RNA-seq, and ONT direct RNA long-read RNA-seq. Additionally, pooled 10x Chromium 3’ single-cell RNA-seq (scRNA) and single-nucleus RNA-seq (snRNA) libraries were available, sequenced with both Illumina short reads and PacBio Kinnex/ONT long reads. Cell line-specific barcode lists for the pooled 10x libraries were obtained from the LongBench project (Ritchie et al., 2025), which assigned cell line identity via SNP-based genotype demultiplexing. We used these barcode lists to demultiplex reads by cell line. PacBio Kinnex provides 8-16x higher throughput than standard IsoSeq, yielding substantially deeper long-read coverage for ground truth generation.

#### Synthetic bulk data

Synthetic RNA-seq reads with known DoGs were generated using polyester (Frazee et al., 2015). We selected 100 protein-coding genes from GENCODE annotations as DoG genes and 100 as negatives. DoG transcripts were modeled with exponential coverage decay downstream of the TES, with a half-life of 4,919 bp measured from PacBio IsoSeq read endpoints in HepG2. DoG lengths varied across genes (5, 10, 20, 50, 100 kb) with readthrough fractions of 10% or 25%. To create an expression sweep, the same 200 genes were used to generate seven separate libraries at gene body coverage levels of 10, 25, 50, 100, 250, 500, and 1000x, with all genes in each library set to the same expression level. Background noise at 1.5% of gene body signal was added to all genes to simulate real intergenic noise.

#### IAV-infected MDM RNA-seq

For read-in gene correction validation (Supplementary Fig S3), we used the influenza A virus (IAV)-infected monocyte-derived macrophage (MDM) RNA-seq dataset from Heinz et al. (2018) (GEO: GSE103477). This is the same dataset used by ARTDeco for read-in gene validation [9]. The dataset includes MDMs infected with wildtype IAV (which causes widespread readthrough via NS1-mediated CPSF30 inactivation), dNS1 mutant IAV (NS1-deleted, reduced readthrough), and mock-infected controls, with two replicates per condition.

Among these datasets, SG-NEx and LongBench are human cancer cell-line samples in which appreciable readthrough is expected; the synthetic dataset provides ground-truth DoGs at controlled levels; and the IAV-infected MDM dataset is a stress-induction model with widespread elevated readthrough used specifically for read-in correction validation. A summary of all datasets used in this study is provided in Table 1.

**Table 1.**
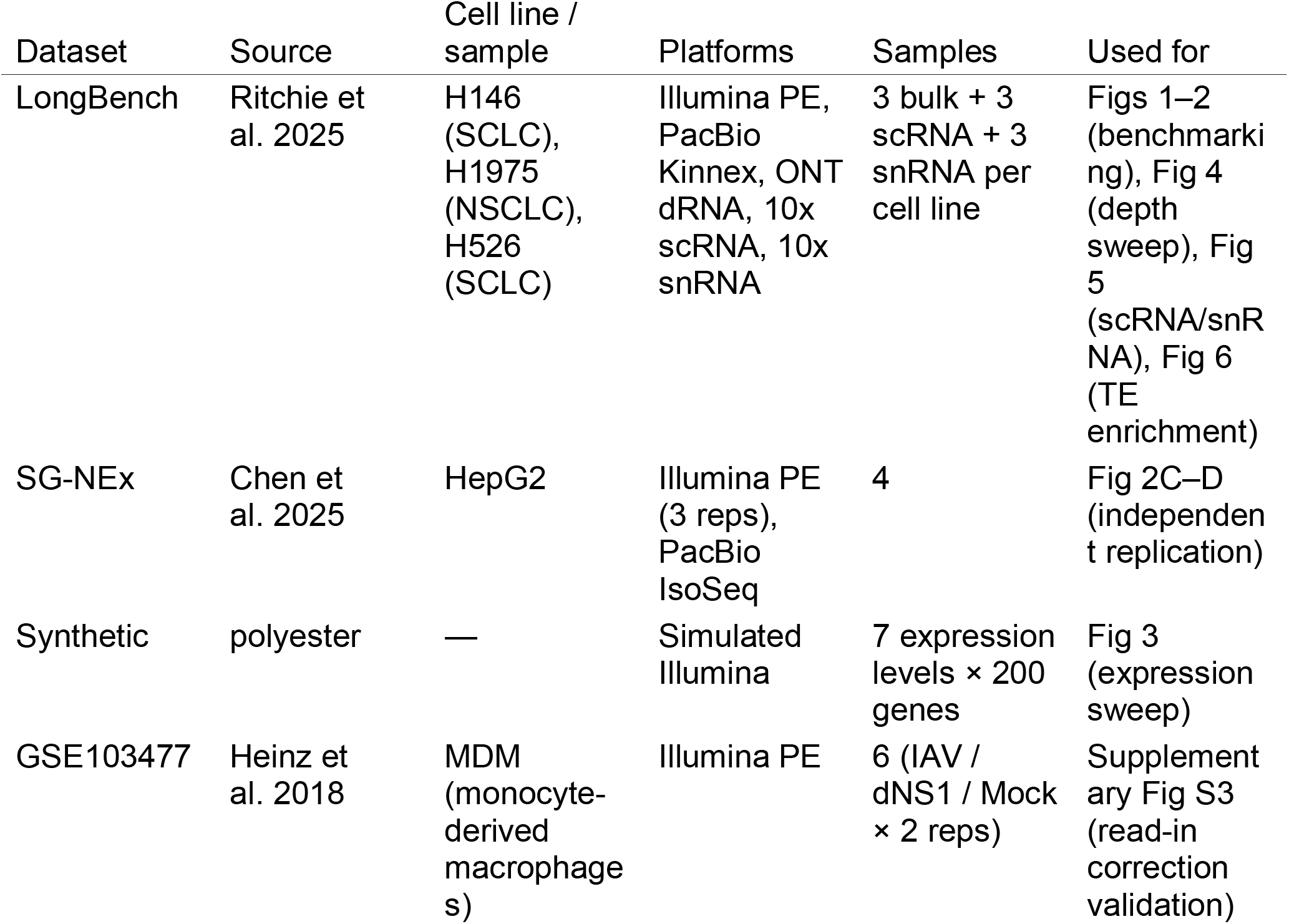
Summary of all datasets used in this study.

### Alignment

Short reads were aligned to hg38 (GENCODE v44) using STAR (Dobin et al., 2013). Long reads were aligned with minimap2 (Li, 2018) using splice:hq for PacBio and splice for ONT. 10x Chromium data was processed with STARsolo for barcode and UMI assignment. Long-read 10x data was demultiplexed using flexiplex for barcode extraction followed by minimap2 alignment.

### Ground truth generation

Long-read ground truth was generated using Dogcatcher’s long-read mode on PacBio Kinnex and ONT BAMs. A 5’ origin filter required each read’s 5’ end to fall within the annotated gene body, preventing misattribution from overlapping downstream genes. Genes with ≥2 readthrough reads extending ≥4 kb past the TES were classified as DoG-positive.

#### PolyA site filtering

DoG regions overlapping known polyadenylation sites were removed from both truth sets and detector calls. GENCODE v46 polyA annotations were used. Reads terminating at downstream polyA sites represent alternative polyadenylation rather than transcription termination defects (Vilborg et al., 2015; Vilborg et al., 2017). Vilborg et al. demonstrated that DoG regions are depleted for canonical AAUAAA polyadenylation motifs compared to non-DoG intergenic regions. Without this filter, 40-75% of apparent DoGs in long-read truth corresponded to alternative polyA usage events rather than genuine readthrough. The polyA filter was applied consistently to both truth and detector calls.

### Benchmarking

Detection performance was evaluated using precision, recall, and F1 score at the pergene level. A true positive is a gene where both the detector and long-read truth identify a DoG ≥4 kb. Boundary accuracy was evaluated using Boundary F1, in which each true positive is weighted by proximity of the called DoG length to the 75th percentile of long-read endpoints for that gene. The 75th percentile was chosen as the truth reference because it represents the predominant termination zone of readthrough transcription. Most polymerase molecules terminate stochastically along the DoG, making the median endpoint reflect early termination events rather than the functional DoG boundary, while the maximum extension is dominated by rare outlier reads. The 75th percentile captures where the bulk of readthrough ends without being skewed by extremes (Supplementary Fig S1). Length ratios (called length divided by truth median, 75th percentile, and max extension) provide additional boundary context.

Dogcatcher detectors were compared against ARTDeco (Roth et al., 2020) run on the same Illumina data with default parameters (500 bp window, 4 kb minimum, 0.15 FPKM threshold).

Sequencing depth sensitivity was assessed by downsampling bulk Illumina BAMs from three LongBench cell lines (H146, H1975, H526) to 75%, 50%, 25%, 10%, 5%, and 1% of total reads and running Dogcatcher2 at each depth against each cell line’s PacBio Kinnex truth.

### Software implementation

Dogcatcher2 is implemented in R with minimal dependencies (GenomicRanges, GenomicAlignments, Rsamtools, rtracklayer, data.table, all standard Bioconductor/CRAN packages commonly available in RNA-seq analysis environments).

No external tools (HOMER, bedops, bowtie2) or cross-language bridges (rpy2) are required, in contrast to ARTDeco which depends on HOMER for tag directory creation, rpy2 for R/Python interoperability, bedops, and RSeQC.

The changepoint package (Killick et al., 2012) is an optional dependency for the PELT detector; the Bayesian changepoint, segmented regression, and sliding window detectors use only base R with no additional packages. The depmixS4 package is an optional dependency for the HMM detector.

Coverage extraction was vectorized using IRanges Views operations, providing substantial speedup over the per-window loop in the original Dogcatcher implementation (Melnick et al., 2019). Roth et al. (2020) reported that ARTDeco was approximately 5-fold faster than the original Dogcatcher; the current implementation narrows this gap to approximately 4.5-fold while adding gene body normalization and multiple detection algorithms. On LongBench bulk RNA-seq, a single Dogcatcher2 detector (segmented regression or Bayesian changepoint) ran in approximately 2.5 hours per sample (H1975: 153M reads; H526: 118M reads) versus approximately 30 minutes for ARTDeco, on 8 cores and 32 GB RAM.

### TE enrichment analysis

To investigate the relationship between transposable elements and DoG regions, we overlapped DoG coordinates with RepeatMasker hg38 TE annotations (5.7 million loci). For each DoG, a matched-length control region was created downstream of a protein-coding gene without DoGs, clipped to the available intergenic space. TE density (elements per kb) was compared between DoG and control regions using Wilcoxon rank-sum tests with Bonferroni correction. Inverted Alu pair annotations were obtained from precomputed dsRNA pair analysis. TE locus-level expression was quantified using LocusMasterTE, which extends the Telescope EM framework with long-read informed priors. Telescope assigns multimapping short reads to TE loci using an Expectation-Maximization algorithm that iteratively updates the posterior probability of each read-locus assignment. LocusMasterTE improves on this by incorporating per-locus expression estimates derived from PacBio Kinnex long-read data as Bayesian priors in the EM, weighting the reassignment toward loci with confirmed long-read expression. This is particularly important for young, highly similar TE subfamilies (e.g. L1HS, AluY) where short-read mapping alone cannot distinguish between copies. The TE annotation GTF used was retro.hg38.v1 (curated HERV and full-length L1 loci, ∼28,000 elements). For single-cell TE quantification, Stellarscope was used in pseudobulk mode on per-cell-line split BAMs from 10x Chromium scRNA-seq and snRNA-seq data.

### Software and data availability

Dogcatcher2 is available at https://github.com/Senorelegans/Dogcatcher2. SG-NEx data is available at s3://sg-nex-data. LongBench data is available at s3://longbench-data.

## Results

### Dogcatcher2 identifies readthrough transcription validated by long-read ground truth (Fig 1)

To illustrate Dogcatcher2’s approach, we examined LINC02697 in H1975, a gene with validated readthrough transcription confirmed by PacBio Kinnex long reads (Fig 1A). Short-read Illumina coverage extends beyond the transcription end site (TES), and individual PacBio long reads independently confirm readthrough, with the 75th percentile of read extensions reaching 11.2 kb. Dogcatcher2’s segmented regression and Bayesian changepoint detectors both identify this DoG from the short-read data alone, while ARTDeco, PELT, HMM, and Sliding Window fail to detect it.

**Fig 1.**
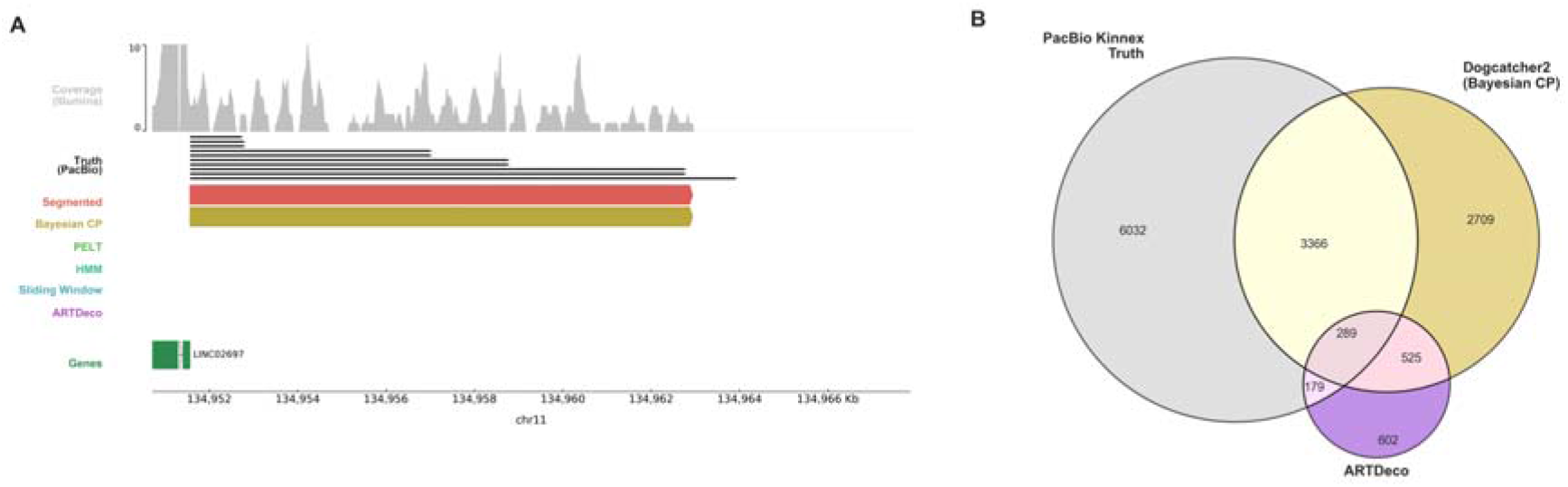
Dogcatcher2 detects readthrough transcription validated by long-read ground truth. (A) Genome browser view of a DoG at LINC02697 in H1975. Top: short-read Illumina coverage (capped at 10x). Individual PacBio Kinnex long reads (grey bars, sampled to 10) show readthrough extending past the TES. Dogcatcher2’s segmented regression and Bayesian changepoint detectors (red, gold) identify the DoG, while PELT, HMM, Sliding Window, and ARTDeco fail to detect it. Gene annotation shown at bottom (+ strand only). (B) Three-way Venn diagram comparing DoG detection in H1975: PacBio Kinnex truth (9,866 genes with validated DoGs ≥4 kb), Dogcatcher2 segmented regression (6,663 called DoGs), and ARTDeco (1,595 called DoGs). Dogcatcher2 recovers 3,429 truth DoGs compared to 468 for ARTDeco, a 7-fold improvement. Pooled results across all three LongBench cell lines shown in Supplementary Fig S2.

Genome-wide, Dogcatcher2’s segmented regression detector identifies substantially more validated DoGs than ARTDeco (Fig 1B). In H1975, PacBio Kinnex long reads establish ground truth for 9,866 genes with readthrough extensions ≥4 kb. Dogcatcher2 recovers 3,429 of these truth DoGs (35% recall), compared to just 468 for ARTDeco (5% recall), a 7-fold improvement. Per-gene Venn assignments are provided in S1 Table. This pattern was consistent across all three LongBench cell lines (Supplementary Fig S2; per-gene data in S2 Table).

### Dogcatcher2 statistical detectors outperform existing methods (Fig 2)

We benchmarked all five Dogcatcher2 detectors and ARTDeco against long-read ground truth from three LongBench cell lines (H146, H1975, H526), using PacBio Kinnex truth with polyA site filtering and a minimum exon coverage of 3x (Fig 2A). Segmented regression achieved the highest F1 (0.334), followed closely by Bayesian changepoint (0.331). Both substantially outperformed ARTDeco (F1 = 0.080), representing an approximately 4-fold improvement. The HMM detector achieved the highest precision (33%) but lower recall, while the sliding window detector showed the lowest performance among Dogcatcher2 methods.

**Fig 2.**
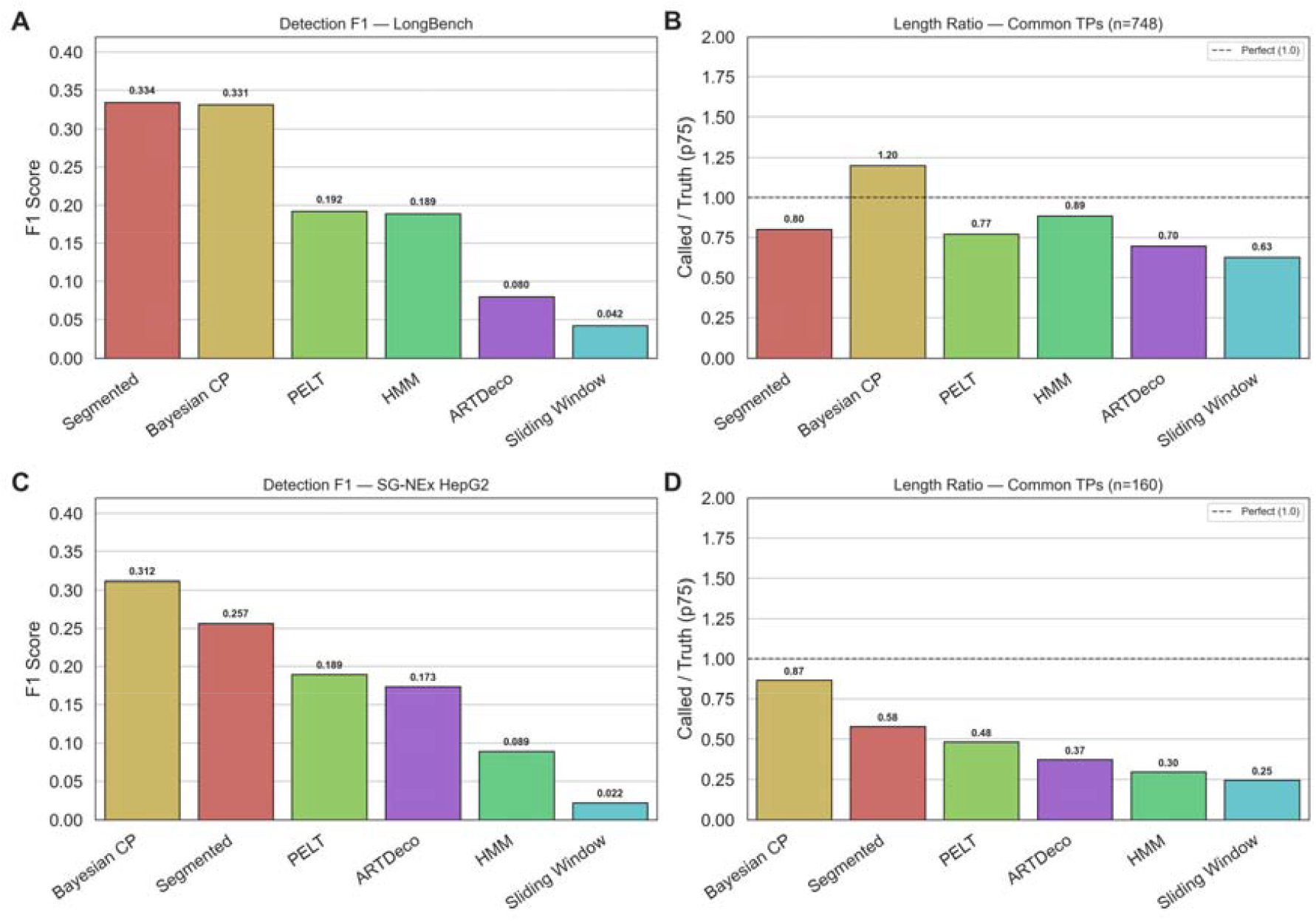
Dogcatcher2 statistical detectors outperform ARTDeco on real data. (A-B) LongBench bulk RNA-seq from three cell lines (H146, H1975, H526) benchmarked against PacBio Kinnex long-read truth with polyA site filtering. (A) Detection F1 score. Segmented regression (0.334) and Bayesian changepoint (0.331) achieve approximately 4-fold higher F1 than ARTDeco (0.080). (B) Length ratio on the 748 common true positives detected by all methods. Called DoG length divided by truth 75th percentile extension. Dashed line indicates perfect agreement (1.0). Bayesian changepoint (1.20) is closest to truth while ARTDeco underestimates (0.70). (C-D) Independent replication on SG-NEx HepG2 with PacBio IsoSeq truth. (C) Detection F1. Bayesian changepoint (0.312) and Segmented (0.257) outperform ARTDeco (0.173). (D) Length ratio on 160 common true positives. Bayesian changepoint (0.87) is again closest to 1.0, while ARTDeco (0.37) substantially underestimates DoG length. Rankings are consistent across both datasets and long-read platforms (PacBio Kinnex vs PacBio IsoSeq).

To fairly compare boundary accuracy we computed length ratios on the 748 genes detected by all three methods (Segmented, Bayesian changepoint and ARTDeco) across all cell lines (Fig 2B). On these common true positives ARTDeco underestimated DoG length the most (ratio 0.70) while Bayesian changepoint was closest to the true boundary (ratio 1.20) and segmented regression slightly underestimated (ratio 0.80). The apparently favorable ARTDeco length ratio on its own detections is partly an artifact of selection bias as ARTDeco detects far fewer DoGs (5.9% recall vs 37% for Dogcatcher2) and its true positives are biased toward high-signal genes where boundaries happen to be clearest.

Performance rankings were consistent across all three cell lines and across truth platforms (PacBio, ONT, and PacBio-ONT consensus), indicating that the results are robust to cell line-specific biology and long-read technology differences. These findings were replicated on the independent SG-NEx HepG2 dataset using PacBio IsoSeq truth (Fig 2C-D), where Bayesian changepoint (F1 = 0.312) and segmented regression (F1 = 0.257) again outperformed ARTDeco (F1 = 0.173), confirming generalization across datasets and long-read platforms. We further validated Dogcatcher2’s read-in gene correction on the same IAV-infected MDM dataset used by ARTDeco (GSE103477), where the correction removed 3 spurious DE genes caused by upstream readthrough contamination, comparable to ARTDeco’s findings (Supplementary Fig S3).

### Synthetic expression sweep reveals detection limits (Fig 3)

To evaluate detection under controlled conditions, we generated synthetic RNA-seq data with known DoGs using polyester [13] (Fig 3). The same 100 DoG genes and 100 negative genes were used across seven expression levels (10, 25, 50, 100, 250, 500, and 1000x gene body coverage), producing separate libraries at each level. Synthetic DoGs were modeled with exponential coverage decay (half-life 4,919 bp, measured from PacBio IsoSeq read endpoints in HepG2) across a range of lengths (5-100 kb) and readthrough fractions (10%, 25%).

**Fig 3.**
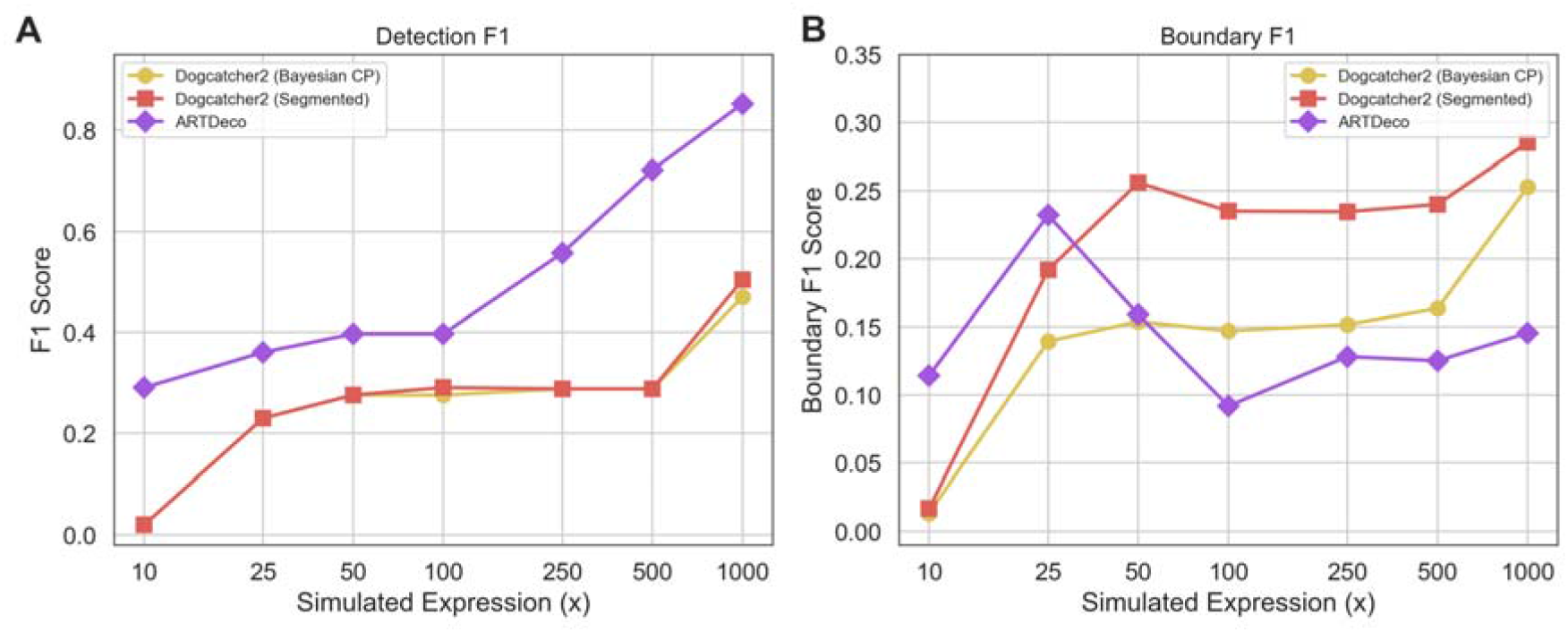
Synthetic expression sweep with decay model. Same 100 DoG genes and 100 negatives evaluated at seven expression levels (10-1000x) using polyester-generated reads with exponential coverage decay. (A) Detection F1 versus simulated expression level. ARTDeco outperforms Dogcatcher2 detectors at all expression levels on clean synthetic data, reflecting the idealized conditions that favor absolute FPKM thresholds. (B) Boundary F1 versus simulated expression level. Segmented regression achieves the best boundary accuracy at moderate expression levels (25-500x), while ARTDeco’s boundary accuracy declines at higher expression despite detecting more DoGs.

ARTDeco outperformed Dogcatcher2 detectors on detection F1 across all expression levels on synthetic data, reaching F1 = 0.85 at 1000x compared to 0.50 for segmented regression and 0.47 for Bayesian changepoint (Fig 3A). ARTDeco’s detection advantage increased with expression level, while Dogcatcher2 detectors showed a more gradual improvement from 25x onward.

However, boundary accuracy told a different story (Fig 3B). Segmented regression achieved the best Boundary F1 across moderate expression levels (25-500x), peaking at 0.26 at 50x, while ARTDeco’s boundary accuracy peaked at 0.23 at 25x and declined at higher expression despite detecting more DoGs. At 1000x, segmented regression (Boundary F1 = 0.29) substantially outperformed ARTDeco (0.15), indicating that ARTDeco detects DoGs but calls their boundaries less accurately as signal increases.

The synthetic data represents an idealized best-case scenario for absolute-threshold methods like ARTDeco. The simulation provides uniform coverage per gene with a clean exponential decay model, minimal background noise (1.5% of gene body), no overlapping genes (≥120 kb intergenic space), and no batch effects or library complexity variation. Under these conditions, ARTDeco’s 0.15 FPKM threshold reliably distinguishes signal from background. Real RNA-seq data violates all of these assumptions where gene coverage is uneven across exons, intergenic regions contain variable noise from DNA contamination and repetitive elements, neighboring genes produce overlapping signal, and expression levels span orders of magnitude within a single library. These factors explain why ARTDeco’s strong synthetic performance (Fig 3A) does not translate to real data, where Dogcatcher2 achieves a 4-fold F1 improvement (Fig 2A). Gene body normalization is robust to these real-world sources of variability because it asks whether downstream signal is meaningful relative to the gene itself, rather than relative to an absolute threshold.

These results establish a practical detection floor: at expression levels below ∼25x, all coverage-based methods struggle to detect readthrough. This is consistent with the depth sensitivity reported for DoGFinder [8] and motivates the min_exon_cov filter in Dogcatcher2.

### Sequencing depth determines detection sensitivity (Fig 4)

The synthetic expression sweep (Fig 3) isolates per-gene signal strength as a variable: all genes in each synthetic library are set to the same coverage, revealing the detection floor for individual genes. In contrast, downsampling real data varies the total library size, where all genes lose signal proportionally and low-expression genes fall below detection thresholds first. To quantify this real-world relationship between sequencing depth and DoG detection, we downsampled bulk Illumina BAMs from three LongBench cell lines (H146, H1975, H526) to 1%, 5%, 10%, 25%, 50%, and 75% of total reads and ran Dogcatcher2 (Bayesian changepoint and segmented) and ARTDeco at each depth against each cell line’s PacBio Kinnex truth (Fig 4).

**Fig 4.**
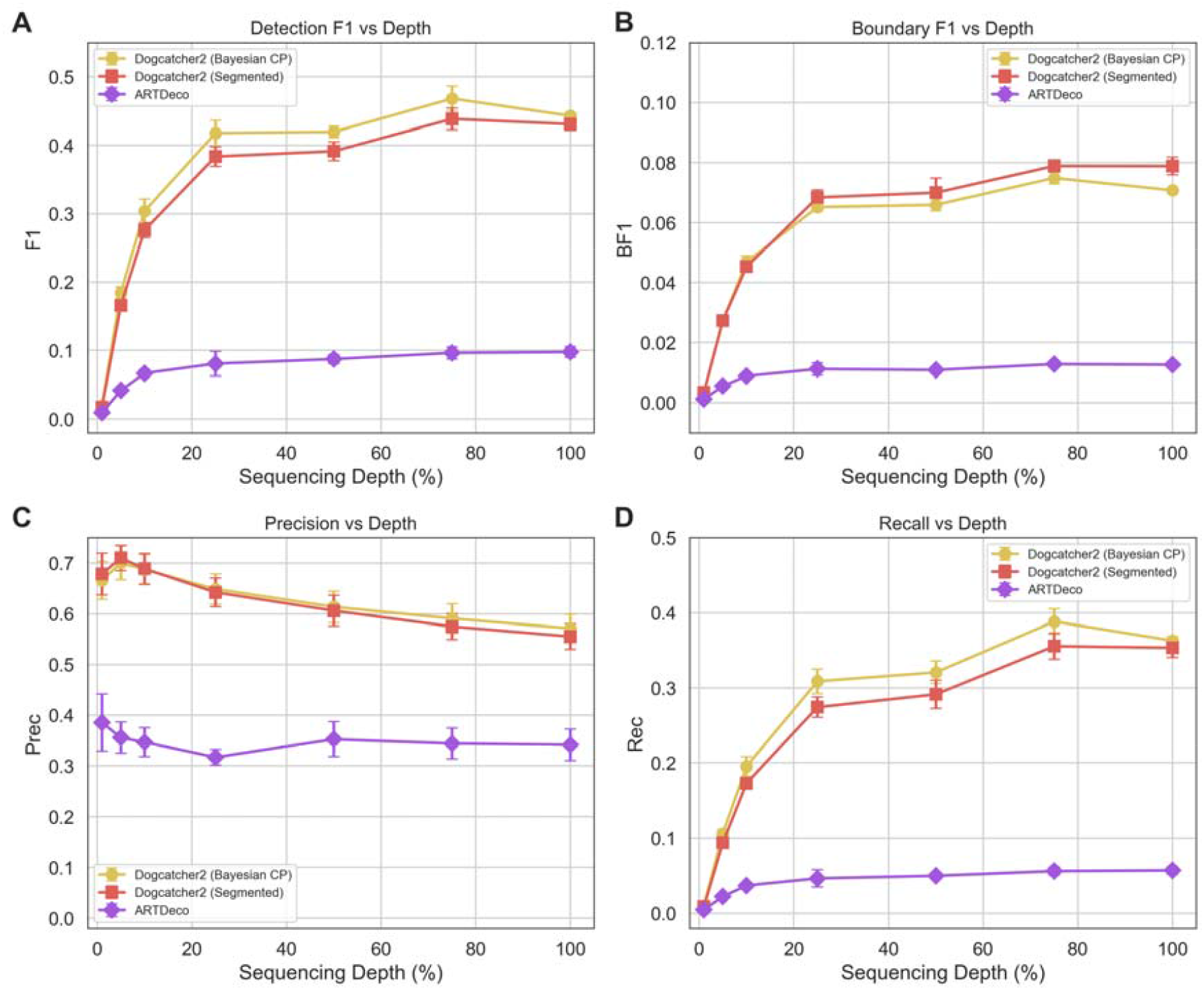
Sequencing depth sensitivity. Bulk Illumina BAMs from three LongBench cell lines (H146, H1975, H526) downsampled to 1-100% of total reads. Lines show mean across cell lines; error bars show SEM. (A) Detection F1 versus sequencing depth. Dogcatcher2 (Bayesian CP) reaches diminishing returns around 25% depth. (B) Boundary F1 versus depth. (C) Precision remains above 60% even at 1% depth. (D) Recall increases monotonically with depth.

Averaged across three cell lines, Detection F1 increased monotonically with depth for Dogcatcher2: 0.020 at 1%, 0.184 at 5%, 0.304 at 10%, 0.418 at 25%, 0.420 at 50%, 0.468 at 75%, and 0.443 at 100% (Fig 4A). Returns diminished substantially beyond ∼25% depth. ARTDeco showed much weaker depth dependence, plateauing around F1 = 0.10 at all depths. Precision remained above 60% for Dogcatcher2 even at 1% depth (Fig 4C), indicating that when the detector makes a call, it is reliable and the primary effect of reduced depth is lower recall (Fig 4D). Boundary F1 followed the same pattern (Fig 4B), with Dogcatcher2 reaching 0.07 at full depth compared to ARTDeco’s 0.01.

Dogcatcher2 at just 5% depth (F1 = 0.184) already surpassed ARTDeco at full depth (F1 = 0.098). Normalization ablation (Supplementary Fig S4) shows that even without gene body normalization the segmented and Bayesian detectors still substantially outperform ARTDeco, indicating that the statistical detection methods themselves are the primary source of improvement. Normalization provides an additional boost particularly at lower depths. These results provide practical guidance for experimental design: ∼25% of total reads (∼12M for a typical experiment) captures most detectable DoGs with diminishing returns beyond this point.

### Pseudobulk scRNA/snRNA detection is depth-dependent (Fig 5)

To assess Dogcatcher2’s applicability to single-cell data, we applied both Bayesian changepoint and segmented regression detectors to pseudobulk profiles generated from LongBench 10x Chromium scRNA-seq and snRNA-seq data, and compared against ARTDeco run on the same pseudobulk BAMs (Fig 5).

**Fig 5.**
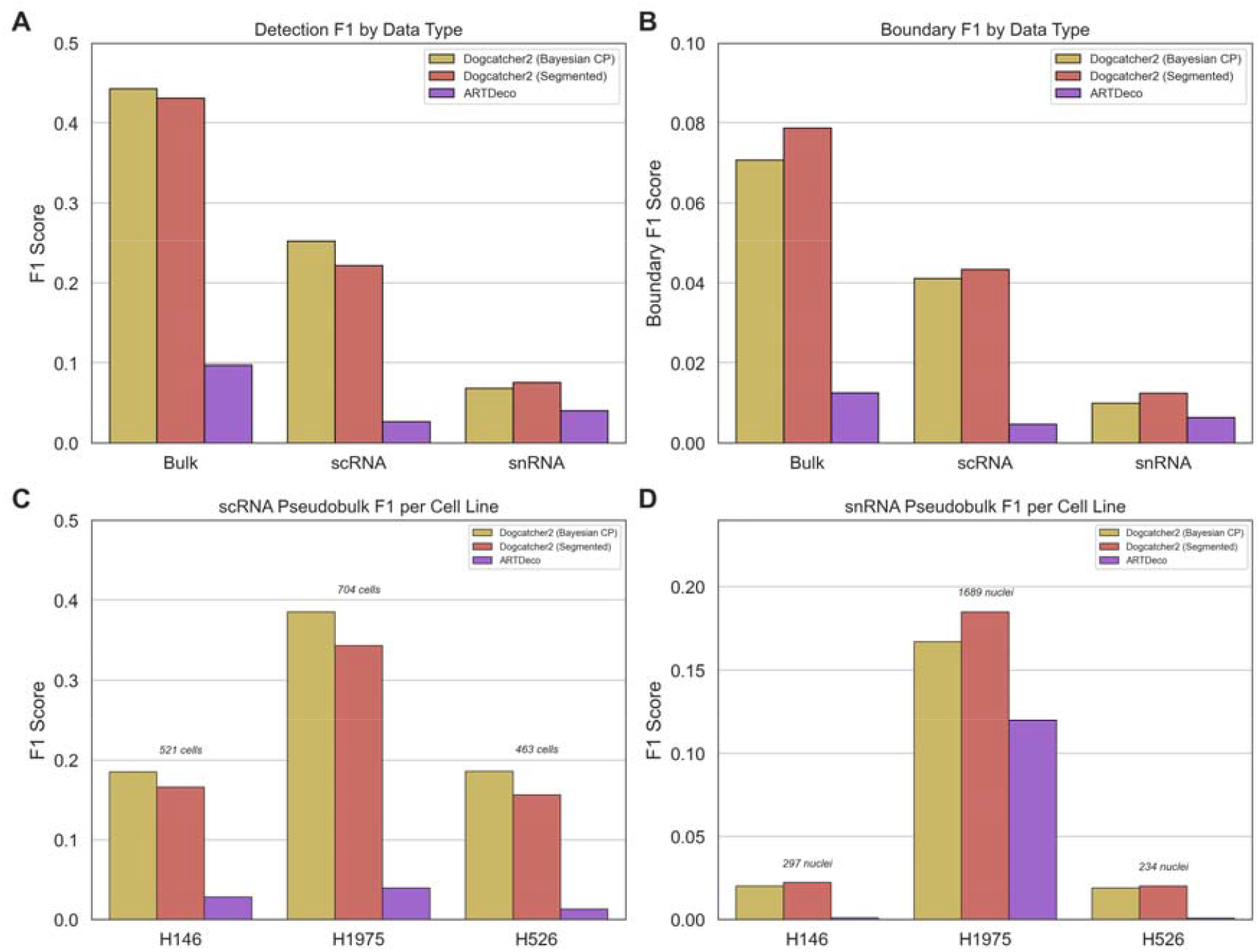
scRNA/snRNA pseudobulk DoG detection. LongBench 10x Chromium data benchmarked against PacBio Kinnex truth. (A) Detection F1 by data type, averaged across three cell lines. Both Dogcatcher2 detectors substantially outperform ARTDeco on all data types. (B) Boundary F1 by data type. (C) scRNA pseudobulk F1 per cell line with cell counts. (D) snRNA pseudobulk F1 per cell line with nuclei counts. Cell/nuclei count is the primary determinant of pseudobulk detection quality.

Both Dogcatcher2 detectors substantially outperformed ARTDeco across all data types (Fig 5A). On bulk data, Bayesian changepoint (F1 = 0.443) and segmented regression (F1 = 0.431) achieved similar performance, both approximately 4.5-fold better than ARTDeco (F1 = 0.098). On scRNA pseudobulk, Bayesian changepoint averaged F1 = 0.252 and segmented regression F1 = 0.222, compared to ARTDeco’s F1 = 0.027. On snRNA pseudobulk, all methods performed lower, with Dogcatcher2 detectors at F1 = 0.069-0.076 versus ARTDeco at F1 = 0.041. Boundary F1 showed the same pattern (Fig 5B), with Dogcatcher2 detectors achieving 2-5x better boundary accuracy than ARTDeco.

Per-cell-line analysis revealed that cell count was the primary determinant of pseudobulk detection quality (Fig 5C-D). For scRNA, H1975 (704 cells) achieved the highest F1 (0.386 Bayesian CP, 0.344 Segmented), while H146 (521 cells) and H526 (463 cells) showed moderate performance. For snRNA, H1975 (1689 nuclei) was the only cell line with meaningful detection (F1 = 0.167-0.185), while H146 (297 nuclei) and H526 (234 nuclei) were near zero. A practical threshold of approximately 500 cells is needed for reliable scRNA DoG detection. The low nuclei counts in these cell lines (H146: 297, H526: 234) likely explain the poor snRNA performance, as pseudobulk coverage at these depths is insufficient for reliable detection.

### DoG regions are enriched for Alu elements and contain actively transcribed TEs (Fig 6)

To investigate the relationship between transposable elements (TEs) and readthrough transcription, we compared TE content in DoG regions against matched-length control regions downstream of non-DoG genes. For each DoG detected by Dogcatcher2 (Bayesian changepoint) in the three LongBench cell lines, we created matched-length control windows downstream of protein-coding genes without DoGs, clipped to the available intergenic space, and counted TE overlaps using full RepeatMasker hg38 annotations.

**Fig 6.**
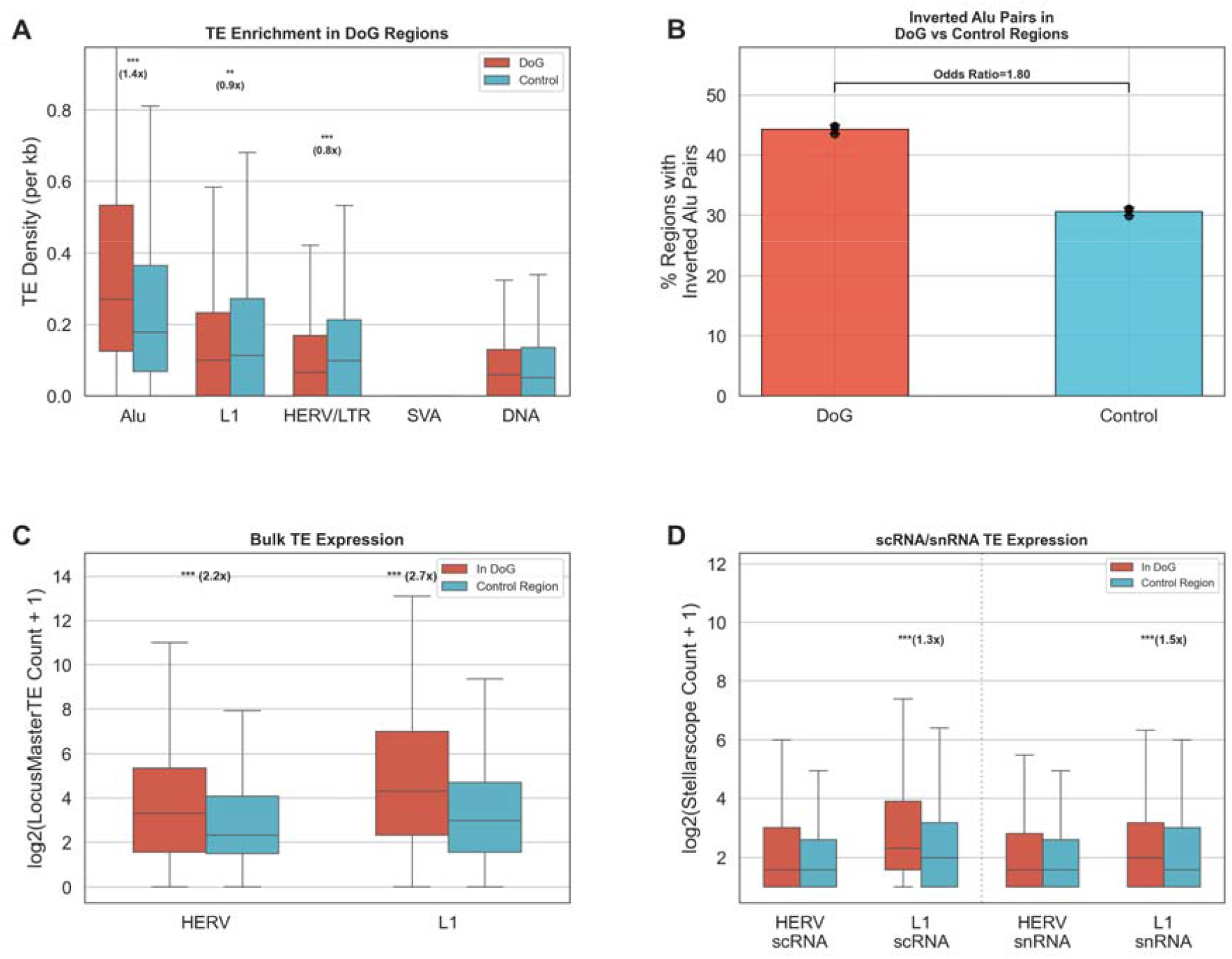
DoG regions are enriched for Alu elements and contain actively transcribed TEs. All data from LongBench, three cell lines (H146, H1975, H526). DoG regions detected by Dogcatcher2 (Bayesian changepoint) compared against matched-length control regions downstream of non-DoG genes. (A) TE density by family. DoG regions are enriched for Alu/SINE elements (1.37-fold, p < 1e-116) but modestly depleted for L1/LINE (0.86-fold) and HERV/LTR (0.77-fold). SVA and DNA transposons show negligible differences. Outliers suppressed for clarity. (B) Fraction of DoG vs control regions containing inverted Alu pairs capable of forming dsRNA. 44.3% of DoG regions contain at least one inverted pair vs 30.6% of controls (Odds Ratio = 1.80). Error bars show SD across three cell lines; dots show individual cell lines. (C) Bulk TE locus-level expression (LocusMasterTE, which integrates PacBio long-read priors with short-read multimapping resolution) for HERV/LTR and L1 elements inside DoG regions vs control regions. TEs within DoGs show 2.2-fold (HERV) and 2.7-fold (L1) higher expression, confirming active transcription through TE insertions. (D) Single-cell TE expression (Stellarscope pseudobulk) for scRNA-seq and snRNA-seq. L1 elements show significant enrichment in DoG regions in both scRNA (1.3-fold, p = 6.1e-16) and snRNA (1.5-fold, p = 1.3e-6). HERV enrichment is not detected at single-cell resolution due to sparse counts. The consistent L1 enrichment across bulk and single-cell platforms validates the readthrough-driven TE expression observed in Panel C.

DoG regions showed significant enrichment for Alu elements (1.37-fold, p < 1e-116) compared to length-matched controls, consistent across all three cell lines (Fig 6A). This enrichment was specific to the Alu/SINE family; L1/LINE elements (0.86-fold) and HERV/LTR elements (0.77-fold) were modestly depleted in DoG regions, while SVA and DNA transposons showed minimal differences.

Given that inverted Alu pairs can form double-stranded RNA (dsRNA) structures that serve as substrates for ADAR editing and activate innate immune signaling through the MDA5/RIG-I pathway, we examined whether DoG regions harbor more dsRNA-forming potential. Using precomputed inverted Alu pair annotations, 44.3% of DoG regions contained at least one inverted Alu pair compared to 30.6% of matched control regions (OR = 1.80, Fig 6B). This suggests that readthrough transcription through Alu-rich downstream regions may generate dsRNA structures with functional consequences for RNA editing and immune activation.

To determine whether TEs within DoG regions are actively transcribed rather than merely present in the underlying DNA sequence, we quantified TE locus-level expression using LocusMasterTE, which integrates PacBio Kinnex long-read priors with short-read multimapping resolution. HERV/LTR loci within DoG regions showed 2.2-fold higher expression than those outside DoGs, and L1 loci showed 2.7-fold higher expression (p = 7.3e-65, Fig 6C). This confirms that readthrough polymerase traverses and transcribes TE insertions in the downstream region.

To validate TE expression enrichment at single-cell resolution, we quantified TE locus-level expression using Stellarscope on pseudobulk scRNA-seq and snRNA-seq profiles from the same LongBench cell lines. L1 elements showed consistent enrichment in DoG regions in both scRNA (1.3-fold, p = 6.1e-16) and snRNA (1.5-fold, p = 1.3e-6) (Fig 6D). HERV enrichment was not detectable at single-cell resolution, likely because HERV elements are longer (5-10 kb) and rarer (∼15,000 annotated loci vs ∼13,000 L1) with fewer reads per locus in sparse single-cell data. The persistence of L1 enrichment across bulk (LocusMasterTE, Fig 6C) and single-cell (Stellarscope, Fig 6D) platforms confirms that readthrough-driven TE transcription is not an artifact of bulk averaging.

While Dogcatcher2 recovers 35% of long-read-confirmed DoGs (7-fold more than ARTDeco’s 5%), a natural concern is whether the remaining undetected DoGs are missed due to repetitive element-driven multimapping artifacts, where short reads fail to map in TE-dense regions, causing coverage dropout that prevents DoG detection. To test this directly, we realigned all three LongBench cell lines with STAR using outFilterMultimapNmax=200 (versus the default of 10) and re-ran Dogcatcher2. Increasing multimapping tolerance had minimal effect on DoG detection: across three cell lines, only ∼230 additional long-read-confirmed DoGs were recovered, and overall recall did not improve (Supplementary Fig S5). This demonstrates that the detection gap is driven by insufficient expression depth at most genes, not by multimapping artifacts in TE-dense regions. Consistent with this, Alu density measured from long-read DoG coordinates was similar across all detection categories, including DoGs found only by long reads (Supplementary Fig S6), indicating that short-read detectors are not systematically biased toward or against TE-dense regions. The Alu enrichment in DoG regions is a genuine biological feature of readthrough-prone loci, not a confound of the detection method.

## Discussion

Dogcatcher2 addresses limitations in existing DoG detection methods by replacing simple threshold-based approaches with statistical detectors operating on gene body-normalized coverage. The central advance is the detector suite itself: segmented regression, Bayesian changepoint, PELT changepoint, and HMM each bring a different statistical framework to the problem of distinguishing readthrough signal from background noise. Previous tools relied on binary coverage thresholds (DoGFinder) or absolute FPKM cutoffs (ARTDeco), which are inherently depth-dependent and do not model the shape of readthrough signal. Gene body exon normalization complements the detectors by making the input signal expression-level-independent: ARTDeco’s 0.15 FPKM threshold treats a weakly-expressed gene’s genuine readthrough the same as background noise, while DoGFinder’s binary 60% coverage criterion requires depth-matched libraries for valid cross-sample comparison [8,9]. Together the normalized signal and statistical detectors ask a different question: does the downstream coverage profile show a statistically significant transition from readthrough to background, relative to the gene’s own expression?

Based on consistent performance across both LongBench and SG-NEx datasets, we recommend Bayesian changepoint as the default detector. While segmented regression achieves marginally higher detection F1 on LongBench (0.334 vs 0.331), Bayesian changepoint consistently produces the most accurate DoG boundaries (length ratio closest to 1.0 on both datasets) and generalizes better across platforms (Fig 2). Bayesian changepoint is set as the default in the Dogcatcher2 software.

The performance advantage of segmented regression and Bayesian changepoint over simpler detectors reflects the biology of readthrough transcription. DoGs exhibit exponential coverage decay downstream of the TES, with a half-life of approximately 5 kb measured from PacBio IsoSeq read endpoints. Segmented regression explicitly models this decay as a piecewise linear function, fitting the transition from active transcription to background. Bayesian changepoint achieves the most accurate DoG boundaries (length ratio 1.20, closest to the ideal 1.0), while segmented regression tends to slightly overestimate DoG length (ratio 1.60). The PELT changepoint algorithm, which assumes piecewise-constant signal, does not capture the gradual decay shape and performs slightly worse for boundary calling despite strong binary detection.

The long-read mode and associated ground truth pipeline fill an important gap. No previous DoG detection tool has been validated against read-level transcript boundaries. Our benchmarks revealed two critical issues in naive ground truth generation: misattribution from overlapping genes (resolved by the 5’ origin filter, which removed 22-35% of false assignments) and confusion with alternative polyadenylation (resolved by the polyA site filter, which removed 40-75% of apparent DoGs that actually represented known polyA usage). These filters are essential for meaningful benchmarking and suggest that previous estimates of readthrough prevalence may include substantial contributions from alt-polyA events.

Read-in gene detection and expression correction address a downstream consequence of readthrough that can confound gene expression analysis. ARTDeco demonstrated that read-in genes produce spurious GO enrichments and eQTLs mapping to upstream DoG-producing genes rather than to the read-in genes themselves [9]. Dogcatcher2 integrates read-in detection directly with DoG calling and provides a correction step in the DESeq2 pipeline to help account for this contamination. We validated this correction on the same IAV-infected MDM dataset used by ARTDeco (GSE103477, Heinz et al. 2018), where IAV causes widespread readthrough via NS1-mediated CPSF30 inactivation. Dogcatcher2’s read-in correction removed 3 spurious DE genes in IAV vs Mock that were falsely called due to upstream readthrough contaminating their counts (Supplementary Fig S3). While the number of corrected genes is small, consistent with ARTDeco’s findings on this dataset, even a few false positives can mislead downstream pathway enrichment and eQTL analyses.

Our depth sensitivity analysis provides practical guidance for experimental design. For bulk RNA-seq, approximately 25M reads captures most detectable DoGs, with diminishing returns beyond this point. For pseudobulk analysis of single-cell data, approximately 500 cells per type are needed for reliable detection. Dogcatcher2 at 5% depth (2.5M reads) already outperforms ARTDeco at full depth (50M reads), demonstrating that the detection methods provide benefits across the depth spectrum. snRNA-seq performed worse than scRNA-seq in our benchmarks partly due to low nuclei counts in the LongBench dataset but also because nuclear RNA captures nascent and incompletely processed transcripts that create a noisier background for DoG detection. Wiesel et al. noted that non-polyA-selected libraries reveal more DoGs [8] which is consistent with this observation as snRNA libraries capture a broader range of nuclear transcripts.

The TE enrichment analysis (Fig 6) connects readthrough transcription to a growing body of work on transposable element activation and innate immune signaling. LaRocca et al. demonstrated that readthrough and intron retention are the primary mechanisms driving transposon expression during aging, with TEs lacking functional promoters becoming transcribed as polymerase reads through from upstream genes [14]. Our finding that DoG regions are enriched for Alu elements and that TEs within DoGs show elevated expression provides direct evidence for this mechanism at individual DoG loci. The enrichment of inverted Alu pairs in DoG regions is particularly relevant because these pairs are the primary substrate for ADAR-mediated A-to-I RNA editing [15,16], and unedited Alu-derived dsRNA can activate the MDA5/RIG-I innate immune pathway [17]. Rutkowski et al. showed that HSV-1 infection causes widespread readthrough that generates dsRNA, while the virus actively degrades it to evade immune detection [18,19]. Our results suggest that stress-induced readthrough in uninfected cells could similarly generate immunostimulatory dsRNA through Alu-rich downstream regions. This is relevant to neurodegenerative diseases where both DoGs and dsRNA accumulation have been observed [20]. Importantly, Dogcatcher2’s detection is not biased against TE-dense DoGs: the multimapping analysis (Supplementary Fig S5) shows the detection gap reflects expression depth, not mapping artifacts. To our knowledge, Dogcatcher2 is the first DoG detection tool that integrates TE annotation overlap, dsRNA pair analysis, and TE locus-level expression quantification directly into the readthrough analysis pipeline.

Several limitations should be noted. DoG boundary accuracy is inherently limited by the resolution of coverage-based methods as detector calls correspond to approximately the 75th percentile of long-read endpoints rather than the maximum extension. Genes with fewer than ∼100x gene body coverage are effectively undetectable by any coverage-based method as the downstream signal falls below noise levels. Our long-read ground truth is biased toward polyadenylated transcripts, meaning non-polyadenylated DoGs, which Vilborg et al. showed constitute a substantial fraction of readthrough transcription [1], are not captured. However, this does not affect the core contribution of the work which is the statistical detection framework itself. The segmented regression and Bayesian changepoint detectors are agnostic to whether the decaying downstream signal originates from polyadenylated or non-polyadenylated readthrough and will work on any coverage profile that exhibits the characteristic decay pattern.

Dogcatcher2 is implemented in R with minimal dependencies, requiring no external tools, cross-language bridges, or complex installation procedures. This contrasts with ARTDeco’s dependency on HOMER, bedops, rpy2, and RSeQC, and with DoGFinder’s reliance on samtools and bedtools. The vectorized implementation using IRanges Views operations addresses the ∼5x speed deficit of the original Dogcatcher noted by Roth et al. [9].

## Acknowledgments

We thank Dr. Robin Dowell (University of Colorado, Boulder) for external review and feedback on this manuscript. Computational analyses were performed on the Alpine high-performance computing cluster.

## Author contributions

Marko Melnick: conceptualization, software, writing (original draft). Christopher D. Link: conceptualization, funding acquisition, writing (review and editing).

## Ethics statement

This study did not involve human subjects, animal subjects, or fieldwork. All analyses were performed on publicly available cell-line and cell-line-derived RNA-sequencing datasets (SG-NEx, LongBench, GSE103477) collected under their respective original protocols.

## Supporting Information Captions

**S1 Fig. Choice of truth reference percentile for boundary accuracy benchmarking**. Median length ratio (called DoG length / truth extension) plotted across five long-read endpoint percentiles (p25, p50, p75, p90, max). At the 75th percentile, all three methods (Bayesian CP, Segmented, ARTDeco) converge closest to 1.0, indicating that detector-called boundaries best match the 75th percentile of long-read endpoints. At lower percentiles (p50), detectors overestimate length (ratios 2-3x), while at maximum extension, detectors underestimate (ratios 0.1-0.5). This justifies using the 75th percentile as the truth reference for Boundary F1 calculations throughout the paper.

**S2 Fig. Three-way Venn diagram of DoG detection pooled across all three LongBench bulk cell lines (H146 + H1975 + H526)**. PacBio Kinnex truth identifies 16,617 genes with validated DoGs. Dogcatcher2 (Segmented) detects 10,874 DoGs, of which 7,143 overlap with truth (66% precision). ARTDeco detects 2,912 DoGs with 1,334 overlapping truth (46% precision). 8,932 truth DoGs are missed by both short-read methods, consistent with the depth sensitivity analysis in Fig 4.

**S3 Fig. Read-in gene correction validation on IAV-infected MDM RNA-seq**. Dataset GSE103477 (Heinz et al. 2018), the same dataset used by ARTDeco for validation. IAV infection causes widespread readthrough via NS1 protein inhibiting polyadenylation (CPSF30 inactivation), while dNS1 (NS1-deleted mutant) serves as a control with reduced readthrough. (A) IAV vs Mock: Dogcatcher2’s read-in correction removed 3 spurious DE genes that were falsely called as differentially expressed due to upstream readthrough contaminating their counts. (B) dNS1 vs Mock: minimal readthrough as expected, correction has negligible effect. The small number of corrections is consistent with ARTDeco’s findings, as read-in effects concentrate at a few highly expressed genes near global DoGs, but even a few false positives can mislead pathway enrichment and eQTL interpretation.

**S4 Fig. Normalization ablation analysis across sequencing depths**. Detection F1 (A), Boundary F1 (B), Precision (C), and Recall (D) shown for Dogcatcher2 detectors with and without gene body normalization, compared to ARTDeco, across seven downsampling levels (1-100%). Both Bayesian CP and Segmented detectors outperform ARTDeco even without normalization, demonstrating that the statistical detection methods themselves are the primary source of improvement. Normalization provides an additional 5-15% F1 boost, most pronounced at lower sequencing depths.

**S5 Fig. Effect of multimapping on DoG detection**. Dogcatcher2 (Bayesian changepoint) run on STAR alignments with outFilterMultimapNmax=10 (default) vs outFilterMultimapNmax=200 for all three LongBench cell lines. (A) Multi-200 produced a net gain of only 66-114 DoGs per cell line. (B) Recall vs PacBio Kinnex truth was unchanged or slightly decreased (H146: 30.9% vs 30.2%), demonstrating that the 65% miss rate reflects expression depth limitations, not multimapping artifacts in TE-dense regions.

**S6 Fig. Venn diagram of DoG detection with Alu density annotations**. Three-way comparison of Long Read Truth, Dogcatcher2 (Bayesian CP), and ARTDeco, with median Alu element density (elements per kb) shown for each section. All DoG regions show elevated Alu density compared to matched non-DoG controls (0.18/kb). Long-read-confirmed regions show 0.62-0.75/kb, while short-read-only DoGs show 0.42-0.47/kb, likely reflecting shorter DoG lengths that sample less Alu-rich downstream sequence.

**S1 Table. Per-gene DoG detection matrix for H1975 (LongBench bulk)**. Tab-separated file with one row per gene and four columns: `gene_id`, `in_truth`, `in_segmented`, `in_artdeco` (boolean). Underlies the three-way Venn diagram in Fig 1B. Truth: PacBio Kinnex long-read endpoints with 5’ origin filtering (≥4 kb extension past TES). Segmented: Dogcatcher2 segmented regression detector. ARTDeco: ARTDeco with default parameters. Genes are restricted to protein-coding genes meeting the gene body coverage threshold.

**S2 Table. Per-gene DoG detection matrix pooled across all three LongBench cell lines (H146, H1975, H526)**. Same column structure as S1 Table. A gene is marked True if any of the three cell lines reports a DoG by that method. Underlies Supplementary Fig S2.

## References

1. Vilborg A, Passarelli MC, Yario TA, Tycowski KT, Steitz JA. Widespread inducible transcription downstream of human genes. Molecular Cell. 2015;59: 449–461. doi:10.1016/j.molcel.2015.06.016

2. Vilborg A, Steitz JA. Readthrough transcription: How are DoGs made and what do they do? RNA Biology. 2017;14: 632–636. doi:10.1080/15476286.2016.1149680

3. Vilborg A, Sabath N, Wiesel Y, Nathans J, Levy-Adam F, Yario TA, et al. Comparative analysis reveals genomic features of stress-induced transcriptional readthrough. Proceedings of the National Academy of Sciences. 2017;114: E8362–E8371. doi:10.1073/pnas.1711120114

4. Rosa-Mercado NA, Bhatt DM. It’s a DoG-eat-DoG world—altered transcriptional mechanisms drive downstream-of-gene (DoG) transcript production. Molecular Cell. 2022;82: 1981–1991. doi:10.1016/j.molcel.2022.04.008

5. Muniz L, Lazorthes S, Delmas M, Rispal J, Gerbaud L, Nicolas E. Who let the DoGs out? – biogenesis of stress-induced readthrough transcripts. Trends in Biochemical Sciences. 2021;46: 1014–1025. doi:10.1016/j.tibs.2021.08.003

6. Melnick M, Gonzales P, Cabral J, Allen MA, Dowell RD, Link CD. Heat shock in c. Elegans induces downstream of gene transcription and accumulation of doublestranded RNA. PLoS ONE. 2019;14: e0206715. doi:10.1371/journal.pone.0206715

7. Abe K, Maunze B, Lopez P-A, Xu J, Muhammad N, Yang G-Y, et al. Downstream-of-gene (DoG) transcripts contribute to an imbalance in the cancer cell transcriptome. Science Advances. 2024;10: eadh9613. doi:10.1126/sciadv.adh9613

8. Wiesel Y, Sabath N, Shalgi R. DoGFinder: A software for the discovery and quantification of readthrough transcripts from RNA-seq. BMC Genomics. 2018;19: 1–10. doi:10.1186/s12864-018-4983-4

9. Roth SJ, Heinz S, Benner C. ARTDeco: Automatic readthrough transcription detection. BMC Bioinformatics. 2020;21: 1–11. doi:10.1186/s12859-020-03551-0

10. Killick R, Fearnhead P, Eckley IA. Optimal detection of changepoints with a linear computational cost. Journal of the American Statistical Association. 2012;107: 1590–1598. doi:10.1080/01621459.2012.737745

11. Love MI, Huber W, Anders S. Moderated estimation of fold change and dispersion for RNA-seq data with DESeq2. Genome Biology. 2014;15: 550. doi:10.1186/s13059-014-0550-8

12. Consortium L. Systematic assessment of long-read RNA-seq methods for transcript identification and quantification. Nature Methods. 2024;21: 1–12. doi:10.1038/s41592-024-02298-3

13. Frazee AC, Jaffe AE, Langmead B, Leek JT. Polyester: Simulating RNA-seq datasets with differential transcript expression. Bioinformatics. 2015;31: 2778–2784. doi:10.1093/bioinformatics/btv272

14. LaRocca TJ, Cavalier AN, Bhatt D, et al. A concerted increase in readthrough and intron retention drives transposon expression during aging and senescence. eLife. 2024;13: RP87811. doi:10.7554/eLife.87811

15. Chung H, Calis JJP, Wu X, Sun T, Yu Y, Sarbanes SL, et al. Human ADAR1 prevents endogenous RNA from triggering translational shutdown. Cell. 2018;172: 811–824. doi:10.1016/j.cell.2017.12.038

16. Kim D et al. Inverted alu repeats: Friends or foes in the human transcriptome. Experimental & Molecular Medicine. 2024;56: 1–11. doi:10.1038/s12276-024-01177-3

17. Ahmad S, Muñoz MV, Guo M, et al. Sensing of transposable elements by the antiviral innate immune system. RNA. 2021;27: 735–752. doi:10.1261/rna.078721.121

18. Rutkowski AJ, Erhard F, L’Hernault A, Bonfert T, Schilhabel M, Crump C, et al. Widespread disruption of host transcription termination in HSV-1 infection. Nature Communications. 2015;6: 7126. doi:10.1038/ncomms8126

19. Esclatine A, Taddeo B, Roizman B. The herpes simplex virus host shutoff (vhs) RNase limits accumulation of double stranded RNA in infected cells: Evidence for accelerated decay of duplex RNA. PLOS Pathogens. 2019;15: e1008111. doi:10.1371/journal.ppat.1008111

20. Ravel-Godreuil C, Massart R, et al. Transposable elements as new players in neurodegenerative diseases. FEBS Letters. 2021;595: 2753–2768. doi:10.1002/1873-3468.14205

