## Supplementary figures and images for "Dogcatcher2: Improved statistical detection of transcriptional readthrough and repetitive element analysis across sequencing platforms"

### Supplemental Figure 1

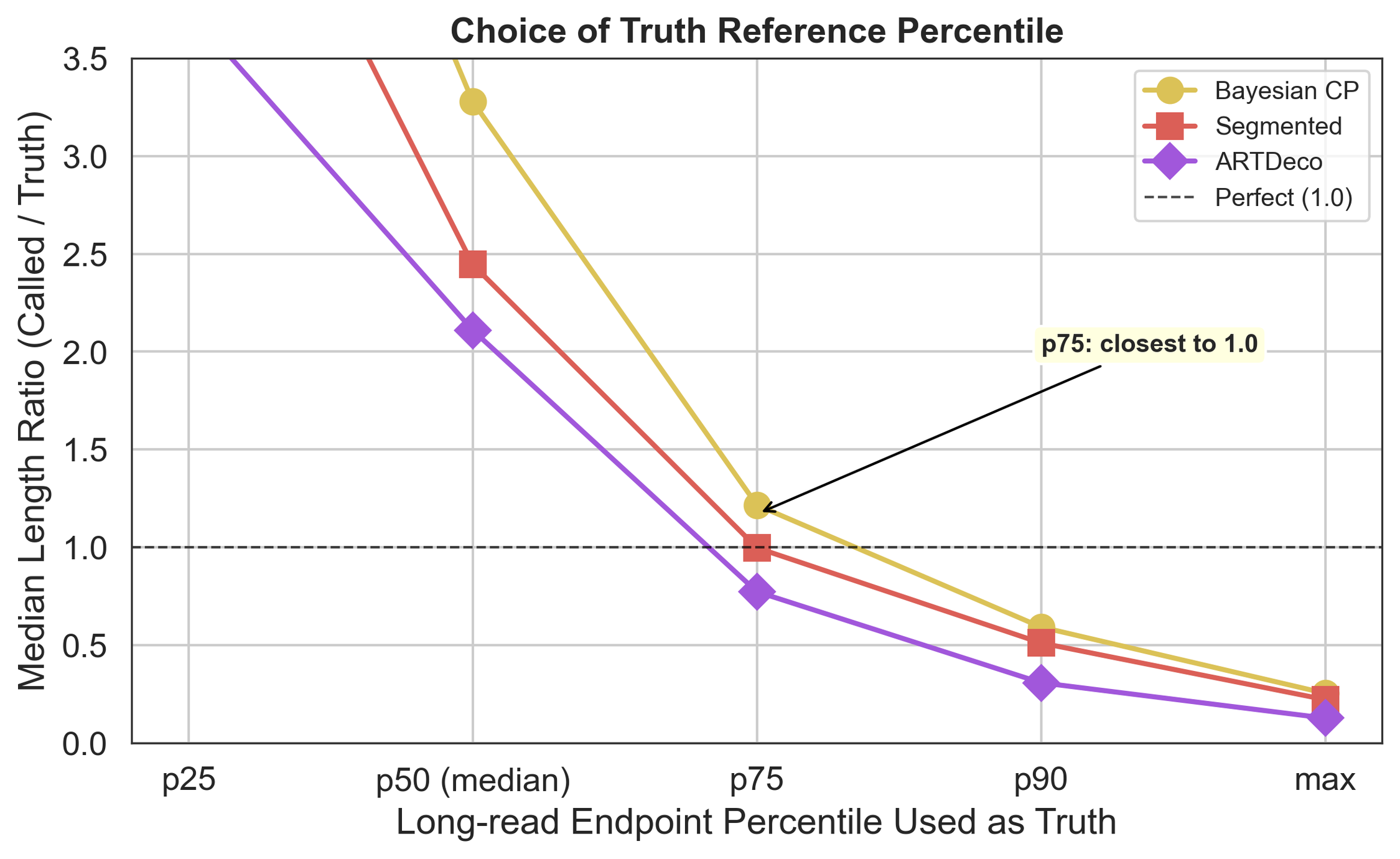

### Supplemental Figure 2

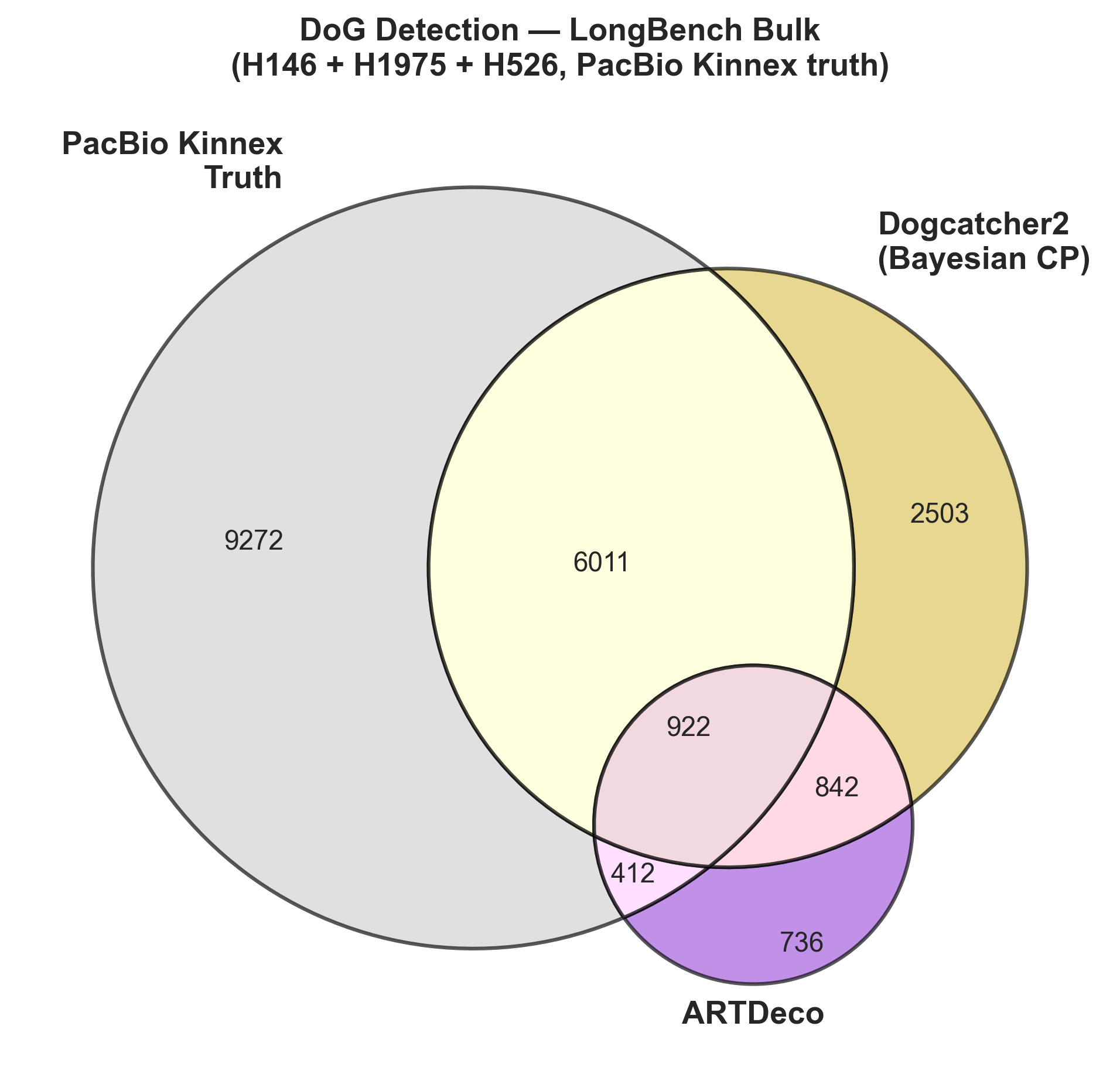

### Supplemental Figure 3

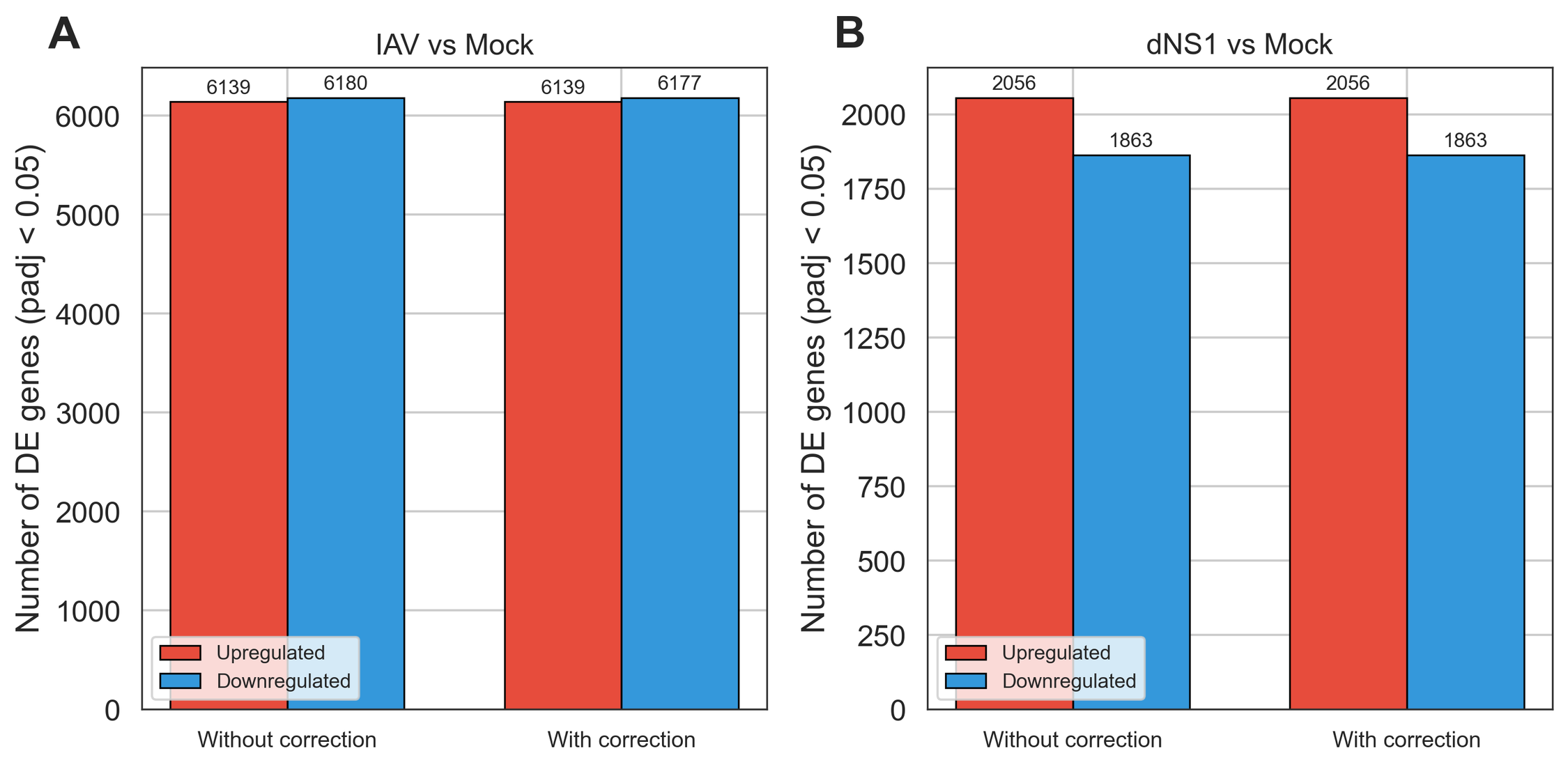

### Supplemental Figure 4

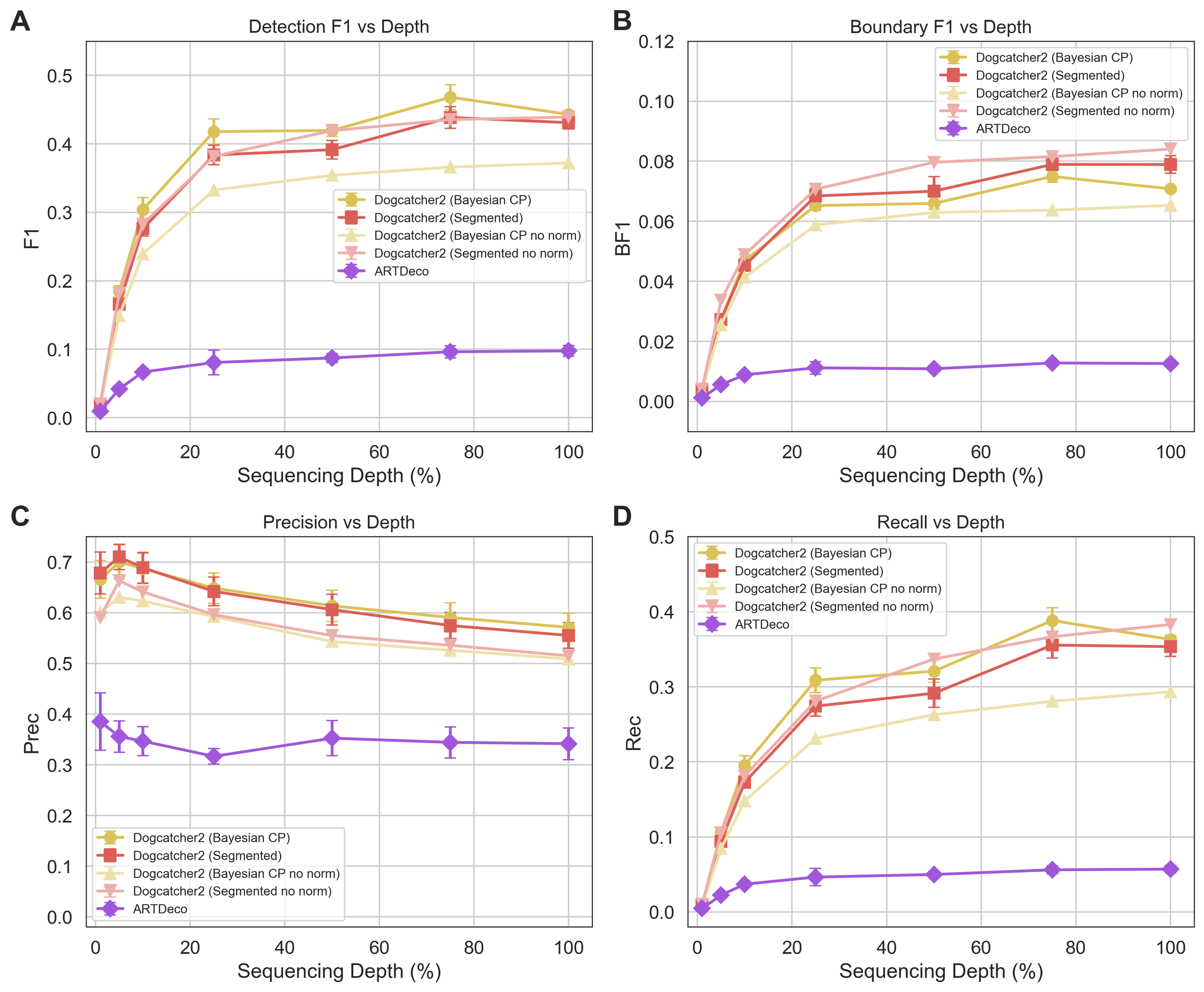

### Supplemental Figure 5

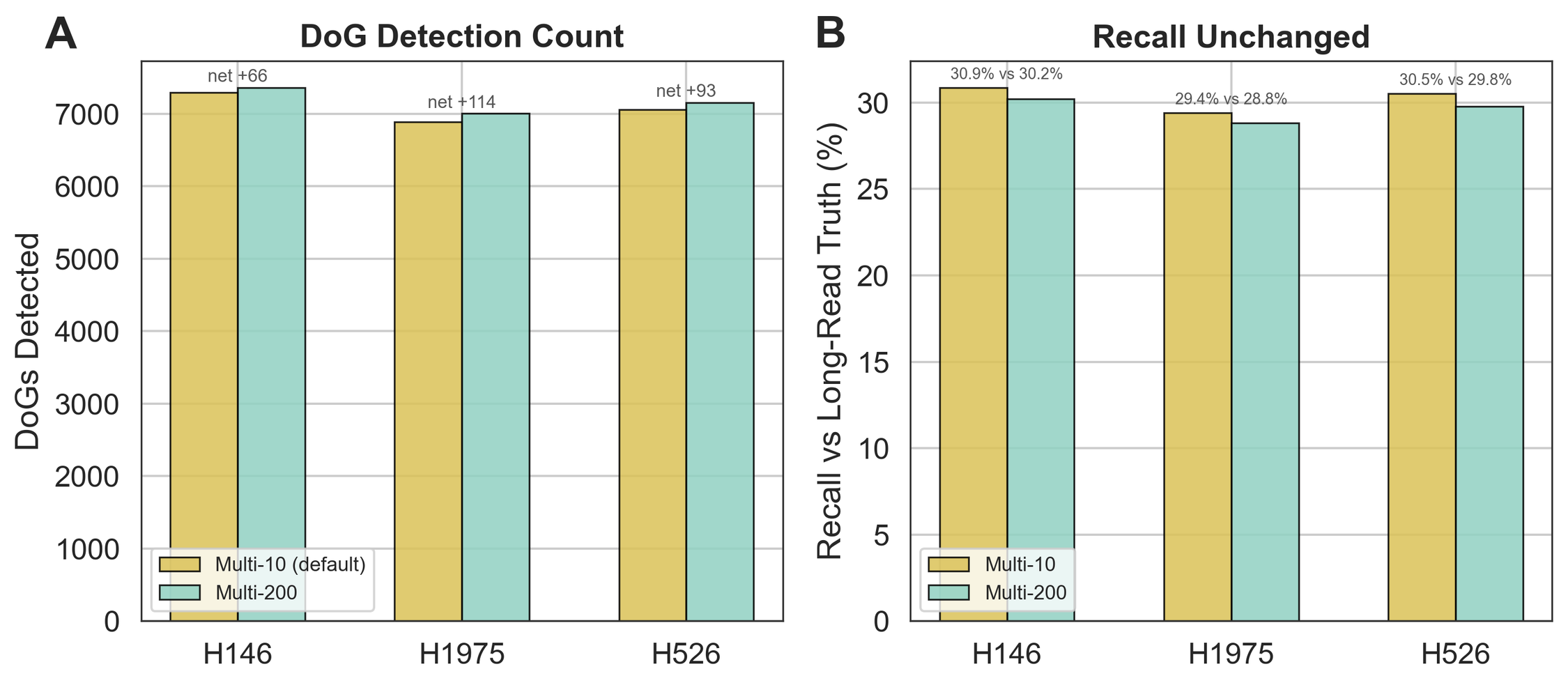

### Supplemental Figure 6

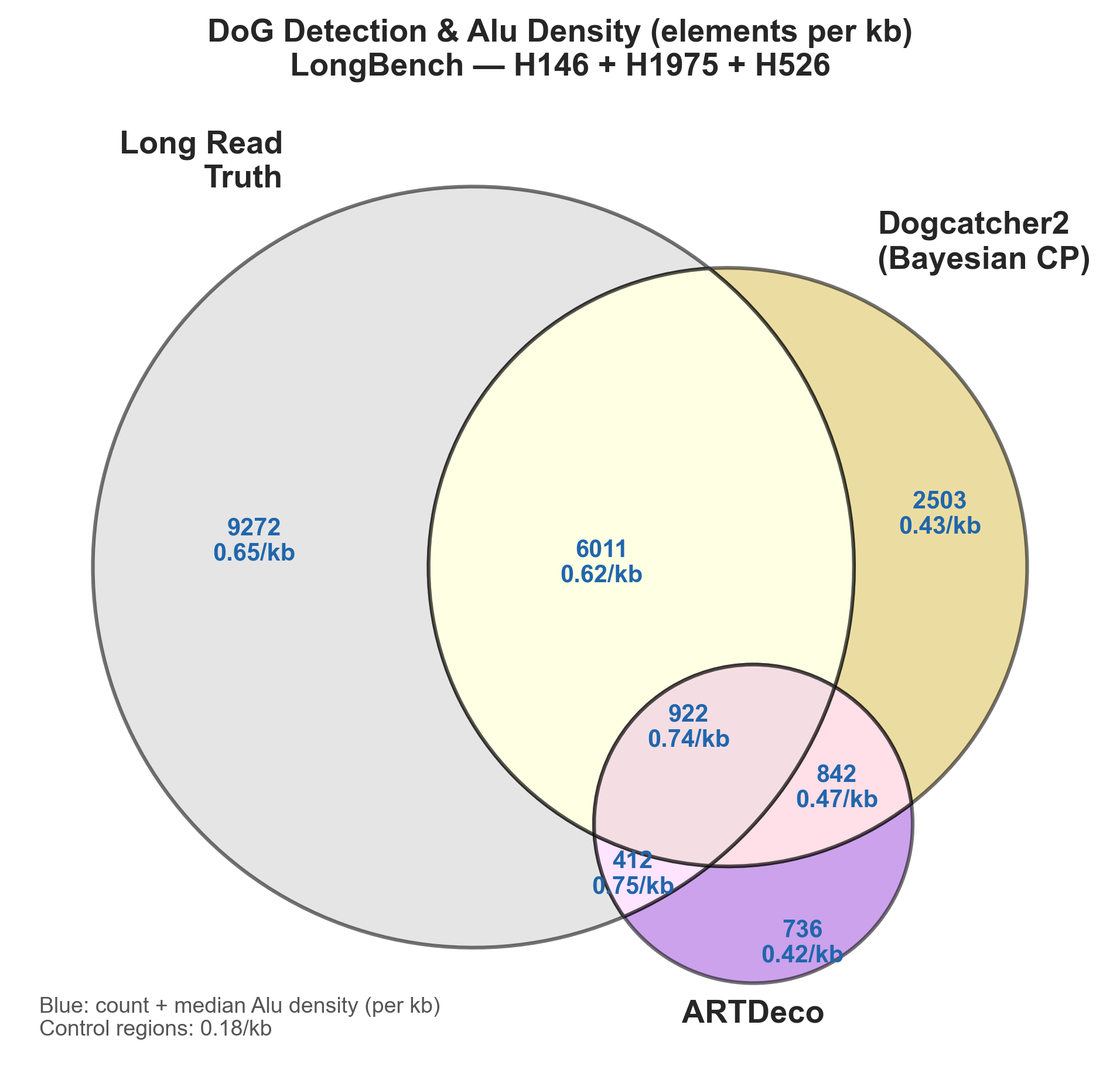
